# Using a residency index to estimate the economic value of saltmarsh provisioning services for commercially important fish species

**DOI:** 10.1101/755835

**Authors:** Hannah A. McCormick, Roberto Salguero-Gómez, Morena Mills, Katrina Davis

## Abstract

Every year, 100 hectares of saltmarsh in the United Kingdom are lost due to sea level rise. The remaining areas are threatened by land conversion, agricultural activities, and climate change. There are important economic consequences to saltmarsh loss, as saltmarsh provides valuable ecosystem services including flood protection, carbon sequestration, and nursery habitat for commercially fished species. Quantifying the economic value of these ecosystem services can help target policies for saltmarsh restoration, or ‘managed realignment’, of new saltmarsh areas. In this study, we quantify the economic value of saltmarsh as a habitat for commercially fished species by developing a residency index. The residency index weights the relative importance of saltmarsh along a species’ lifecycle by explicitly incorporating the target species’ life histories and the estimated proportion of time it spends in saltmarsh at juvenile and adult life stages. Using this index, we estimate the value of saltmarsh to UK commercial fisheries landings. We find that UK saltmarsh contributes annually between 16.7% and 18.2% of total UK commercial landings for European seabass (*Dicentrarchus labrax*), European plaice (*Pleuronectes platessa*), and Common sole (*Solea solea*). Our findings highlight the importance of saltmarsh protection and restoration. Furthermore, our approach provides a general framework that integrates population ecology methods and economic analyses to assess the value of saltmarsh and other coastal habitats for fisheries worldwide.

## INTRODUCTION

Globally, coastal-habitat extent is in severe decline (Waycott et al. 2009; Barbier et al. 2011; Balke et al. 2015). This decline is caused by a variety of human-related activities including climate change, runoff, and coastal development (Waycott et al. 2009; Balke et al. 2015). Rates of decline vary between different coastal habitats. For example, more mangrove forests (35%) have been lost or degraded worldwide, than coral reefs (30%) or seagrass communities (29%; Barbier et al., 2011). Saltmarsh has experienced the most drastic decrease, with 50% of the world’s saltmarshes lost or rapidly declining (Barbier et al., 2011). This decrease is further accentuated by the smaller extent of global saltmarsh compared to other coastal habitats. Currently, saltmarshes occupy 5.5 million ha worldwide (McOwen et al. 2017), compared to 13.8 million ha of mangrove forests (Giri et al 2010), 28.4 million ha of coral reefs (Spalding et al 2001), or 17.7 million ha of seagrass (Green & Short 2003). The rapid rate of decline and comparatively smaller extent of saltmarsh, relative to other coastal habitats, highlights the urgency of prioritising saltmarsh conservation (Colclough et al. 2005).

Saltmarsh and other coastal habitats provide a range of key ecosystem services. These services include flood defence (King & Lestert 1995), recreation (Luisetti et al. 2011), carbon sequestration (Luisetti et al. 2014), and habitat for commercially important fish species (Green et al. 2012). Destruction of coastal habitats like saltmarsh can reduce these ecosystem service flows, leading to large economic losses (Luisetti et al. 2011). For example, Woodward & Wui (2001) estimated that, across the globe, losing one hectare of saltmarsh would result in a loss of $1,212 (USD) per year in recreational birdwatching values and $393 per year in flood defence. Quantifying the economic value of the ecosystem services provided by habitats like saltmarsh allows policy makers to evaluate the impacts of different land use policies, e.g. the economic impacts of habitat restoration (Bateman 2018; Davis et al. 2018). However, government resources for habitat conservation are limited, and so policies to protect coastal habitats, including saltmarsh, must have clear, quantifiable benefits for social welfare (Luisetti et al. 2011; Bateman 2018). To understand whether the societal benefits provided by saltmarsh are greater than the costs of restoring them, it is first necessary to accurately estimate the economic value of the ecosystem services saltmarsh provides (Barbier *et al*., 2011).

One of saltmarsh’s most highly valued ecosystem services is to fisheries (Woodward & Wui 2001). Globally, many commercially important fish species use saltmarsh, including sea mullet (*Mugil cephalus*) in Australia (Taylor et al. 2018) and European seabass (*Dicentrarchus labrax*) in the UK (Colclough et al. 2005). Fisheries value is also one of the most difficult ecosystem services to quantify in saltmarsh (Woodward & Wui 2001). This is because fish will use saltmarsh differently depending on a range of factors including the species, age of fish, time of year, and tidal range (Scott 1999). The same fish species will also use saltmarsh differently in different regions. Tidal range varies in different parts of the world, and this can limit fish access to saltmarsh (Fonseca et al. 2011). Because different species of fish use saltmarsh at different life stages and for different purposes (e.g. feeding, reproduction), a species-specific approach to quantifying the fisheries value of saltmarsh is needed (Scott 1999).

Here, we use a flexible framework to quantify fishery benefits of saltmarsh that integrates economics and population ecology. We use a residency index, as well as expert elicitation and a literature review, to estimate the value of saltmarsh as a habitat for commercially important species of fish. Our approach rests on the assumption that the amount of time a fish spends in saltmarsh (*i*.*e*., residency) is a proxy for its dependency on that habitat. Previously, a residency index methodology has been used to estimate the value of seagrass as a structural habitat for fish (Scott 2000; McArthur & Boland 2006; Jackson et al. 2015). However, to our knowledge, this approach has never been used to estimate the value of saltmarsh. We demonstrate our approach using saltmarsh extent in the UK as a case study. Here, we estimate the value of UK saltmarsh as a habitat for five species of commercial interest (ICES 2018) commonly found in UK saltmarsh (Green et al. 2012). We collect species-specific demographic information from the COMADRE Animal Matrix Database (Salguero-Gómez et al. 2015) and additional resources used to compile the COMADRE matrices (Gerber & Heppell 2004; Hart & Cadrin 2004; Vélez-Espino & Koops 2012). For robustness, we estimate the proportion of time species spend in saltmarsh at different life stages using two independent methods. The first method involves expert elicitation (Scott 2000; McArthur & Boland 2006); and the second a literature review on species’ habitat use (Jackson et al. 2015). We apply our saltmarsh residency index value for each species to catch data from the International Council for the Exploration of the Sea (ICES), and landings data from the Marine Management Organisation (MMO). By applying this index to European landings data, we identify the economic value of UK saltmarsh to UK and European fisheries. Our results provide policy makers and conservation organisations with valuable information to prioritise the restoration of historical saltmarsh sites, and the conservation of existing saltmarsh. The proposed framework, incorporating economics and population ecology, can be used to evaluate the fisheries benefits provided by saltmarsh or other coastal habitats around the world.

## METHODS

### Study site

We focused on the United Kingdom (UK), which has 45,000 ha of saltmarsh (Wolters et al. 2005; Jones et al. 2011), and supports a fleet of 6,238 fishing vessels (Uberoi 2017). The total value of fish landings brought in from the UK fleet in 2015 was £775 million (Marine Management Organisation 2016). This value corresponded to approximately 14.1% of the total landings value of the European fleet in 2015, which was £5.5 billion (Scientific Technical and Economic Committee for Fisheries (STECF) 2017).

### Species identification

We apply our framework to five fish species: *Dicentrachus labrax, Pleuronectus platessa, Solea solea, Chelon labrosus* and *Chelon ramada* (Table 1). We chose these study species based on two criteria: they spend time in UK saltmarsh at some point in their lifecycles (Colclough et al. 2005; Green et al. 2012), and are commercially fished species with available landings data (ICES 2018).

**Table 1.** Life history traits used in this analysis for our five target fish species, including age at maturity, time (in years) spent as a juvenile (t_ji_), time (yr) spent as an adult (t_ai_), natural mortality (m_t_), and number of life stages. (Hart & Cadrin 2004)

| Species Name | Scientific Name | Family | Proxy Species | Proxy Species Family | Age at Maturity (years) | $t_{ji}$ | $t_{ai}$ | $m_{ti}$ | Number of life stages | Reference |
| --- | --- | --- | --- | --- | --- | --- | --- | --- | --- | --- |
| European seabass | *Dicentrarchus labrax* | Moronidae | *Morone saxatilis* | Moronidae | 2.5 | 2.48 | 2.5 | 0.643 | 3 | Velez-Espino & Koops, 2012 |
| European plaice | *Pleuronectus platessa* | Pleuronectidae | *Limanda ferruginea* | Pleuronectidae | 5.52 | 2.35 | 5.52 | 0.665 | 7 | Hart & Cadrin, 2004 |
| Common sole | *Solea solea* | Soleidae | *Limanda ferruginea* | Pleuronectidae | 5.52 | 2.35 | 5.52 | 0.665 | 7 | Hart & Cadrin, 2004 |
| Thicklip grey mullet | *Chelon labrosus* | Mugilidae | *Melanogrammus aeglefinus* | Gadidae | 1.65 | 1.64 | 1.42 | 0.079 | 3 | Gerber & Heppell, 2004 |
| Thinlip grey mullet | *Chelon ramada* | Mugilidae | *Melanogrammus aeglefinus* | Gadidae | 1.65 | 1.64 | 1.42 | 0.71 | 3 | Gerber & Heppell, 2004 |

### Calculating saltmarsh residency indices

Following Scott (1999), we assumed that saltmarsh residency is a proxy for saltmarsh dependency. This allowed us to formulate an index quantifying fish dependency on saltmarsh based on earlier work by Scott (1999):

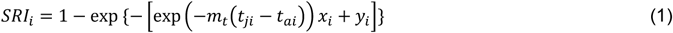

Equation 1 quantifies the Saltmarsh Residency Index (*SRI*) for species *i*, where *m* is the natural (base-line) mortality rate; *t*_*ji*_ is the time (years) spent as a juvenile (*j*) and *t*_*ai*_ as an adult (*a*); *x*_*i*_ is proportion of time spent in saltmarsh as a juvenile; and *y*_*i*_ is proportion of time spent in saltmarsh as an adult (Scott 2000; Jackson et al. 2015). It is important to note that equation 1 takes into account variable habitat use by juvenile and adult life stages (Scott 2000). This distinction is necessary for the SRI of a given species because some fish species, such as *D. labrax*, use saltmarsh primarily as a nursery habitat (Colclough et al. 2005), but can spend the rest/part of their life cycles in other regions. Moreover, the mortality rate *m* of fish often depends on their life stage (Scott 2000; Caswell 2001). Thus, the quantification of the SRI (equation 1) weights time spent in saltmarsh at juvenile and adult life stages based on potential differences in relative mortality rates.

We sourced the demographic data needed to calculate a SRI for each species using matrix population models from the COMADRE Animal Matrix Database (Salguero-Gómez et al. 2015) and additional demographic literature. COMADRE is an open-source demographic database that compiles thousands of matrix population models from hundreds of animal populations published in the peer-reviewed literature. Briefly, a matrix population model incorporates the vital rates (i.e. survival, development, and reproduction) that control the viability and dynamics of a population of interest while explicitly incorporating the contributions of individuals in the population along its lifecycle (Caswell 2001). For the latter, a matrix model includes the aforementioned vital rates for each of the stages in the lifecycle (e.g. juveniles and adult, as necessary for equation 1).

Because not all five exact species are currently available in the COMADRE database, we used matrix population models for closely related species and with similar life history traits (see Table 1), a common practice in comparative fish demography (Vélez-Espino et al. 2006). This approach is based on the fact that demographic rates tend to be well-preserved within derived lineages of the Animal kingdom (Blomberg & Garland 2002, R. Salguero-Gómez, pers. comm. 2018). For *D. labrax* (Moronidae), we used existing COMADRE matrix population models for a closely related species, the striped seabass (*Morone saxatilis*, Moronidae) (Doyle et al. 2017). Because *P. platessa* (Pleuronectidae) and *S. solea* (Soleidae) are also closely related (Hensley 1997), we used matrix population models for a demographically similar species in the COMADRE database from the flatfish family, the yellowtail flounder (*Limanda ferruginea*, Pleuronectidae) (Hart & Cadrin 2004). Similarly, *C. ramada* (Mugilidae) and *C. labrosus* (Mugilidae) are closely related species, and we used matrix population models from a species with similar life history traits, haddock (*Melanogrammus aeglefinus*, Gadidae) (Wright & Tobin 2013).

The demography of each species in COMADRE is archived into associated sub-matrices. These describe vital rates as a function of a variable number of stages, often pre-determined by the authors (Salguero-Gómez et al. 2016). The number of stages in the life cycle also determines the dimensionality of the matrix (Caswell 2001), which can affect model outputs (Enright et al. 1995; Salguero-Gómez & Plotkin 2010). Because the dimensionality of the matrix population models in COMADRE ranges (Salguero-Gómez et al. 2016) (see Table 1), we first collapsed each matrix into a 2×2 model containing a juvenile and an adult stage following the method described by Salguero-Gómez & Plotkin (2010). All initial stages prior to the first reproductive stage were considered juveniles, and all reproductive stages were considered adults. Next, we used methods by Caswell (2001) and Cochran & Ellner (1992) to obtain the age-specific survivorship curve (*I*_*X*_). We used the survivorship curve to calculate age-specific mortality rates (1-*I*_*X*_) and used the mean of the linear mortality models at each stage to represent stage-specific mortality. We calculated natural mortality (*m*_*t*_) rates by averaging these age-specific linear mortality models across the entire lifespan. We also quantified the juvenile residence time (*t*_*ji*_) and adult residence time (*t*_*ai*_) by calculating the fundamental matrix, also following methods by Caswell (2001).

### Estimating proportion of time spent in saltmarsh

There are few quantifiable, habitat-specific demographic studies for fish (Vasconcelos et al. 2014). Consequently, the proportion of time fish species spend in saltmarsh is not readily available in the peer-reviewed or grey literature. To overcome this data shortcoming, we used expert elicitation and conducted a literature review to compile a database of fish habitat usage. These two approaches have previously been used by Scott (1999) and Jackson et al. (2015) to estimate the proportion of time that fish species spend in seagrass in the Mediterranean. We compared the results from both methods with a sensitivity analysis (below).

#### Expert elicitation

Expert opinion is a useful tool for estimating uncertain values (Speirs-Bridge et al. 2010; Hanea et al. 2016; Hemming et al. 2018). We conducted our expert elicitation for the proportion of time our five target species spend in saltmarsh according to recognised standards designed to reduce bias and overconfidence (Hanea et al. 2016). Our process followed key elements of the IDEA protocol outlined in Hemming et al., (2018). The key elements of the IDEA protocol are “investigating” the question, “discussing” as a group, “estimating” final values individually, and “aggregating” the estimations (Hemming et al. 2018). We recruited six experts with UK saltmarsh research or work experience. Each participant received an online questionnaire asking them to estimate the proportion of time our five target fish species spend in saltmarsh during each season, as both a juvenile and an adult (see Appendix S1). The questions were structured by season because many fish species follow migration patterns depending on the time of year (Colclough et al. 2005). When designing the questionnaire we followed the outline by Speirs-Bridge and collaborators (2010), which includes eliciting: minimum, maximum, and best guess value, as well as level of confidence. Best guess estimations were standardised to fit a confidence level of 80% (Hemming et al. 2018). The seasonal estimates of the experts were aggregated to produce an estimate of the proportion of time per year that juveniles and adults of each species spend in saltmarsh.

#### Literature review

The second method for estimating the proportion of time our five target fish species spend in saltmarsh was a meta-analysis of habitat use through a literature review (Jackson et al., 2015). The purpose of the meta-analysis was to determine how many habitats each species uses as a juvenile and as an adult, separately. Due to the lack of habitat-specific demographic information in the literature (Vasconcelos et al. 2014), our approach was based on the assumption that, at a particular life stage, a fish spends equal amounts of time in every habitat it uses at that stage. For example, if a fish can be found in four different habitats as a juvenile, one of which is saltmarsh, we assume that it spends 25% of its time in saltmarsh. This assumption simplifies species’ habitat use, and therefore introduces the possibility of under- or over-estimating the importance of saltmarsh compared to other habitats. We later explore the implications of this assumption through a sensitivity analysis.

We conducted a systematic meta-analysis of habitat use for each study species for all years available in Web of Science and SCOPUS. For full search strings, see Appendix S2. The criteria for inclusion of a peer-reviewed publication was that it must (i) be written in English, (ii) focus on at least one of our five study species, and (iii) provide habitat use information. Because a limited number of studies took place in the UK, we included studies from other regions that had a focus on the species in question and specified habitat use, resulting in a total of 122 sources (See Appendix S2).

To allow for repeatability of habitat classification, habitats were classified according to the European Union Nature Information System (EUNIS) (European Environment Agency 2018). Under the assumption that a fish of a particular life stage uses all habitats equally, habitats were unweighted. If the species was found to spend time in saltmarsh at that life stage, we calculated overall proportion of time spent in saltmarsh as a juvenile and an adult using the following calculations, adapted from Jackson et al. (2015)

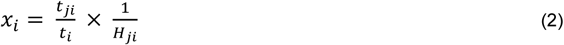

and

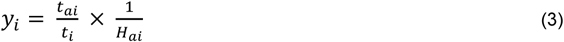

where *x*_*i*_ is proportion of time spent in saltmarsh as a juvenile, *t*_*ji*_ is time spent as a juvenile, *t*_*i*_ is total lifespan, *H*_*ji*_ is total number of habitats used as a juvenile, *y*_*i*_ is proportion of time spent in saltmarsh as an adult, *t*_*ai*_ is time spent as an adult, and *H*_*ai*_ is total number of habitats used as an adult.

### Commercial fisheries landings

To obtain information regarding yearly landings for each species in UK waters, we used ICES time series catch data (ICES 2018). These data identify the species, the ICES area in which the fish were caught, and the live weight in tonnes. We considered catch data from nine ICES fishing areas that border the UK coastline (ICES area codes: 27.4.a, 27.4.b, 27.4.c, 27.7.d, 27.7.e, 27.7.f, 27.7.g, 27.7.a, 27.6.a) (Figure 1). These areas were selected under the assumption that the fish that have spent time in UK saltmarsh are most likely to be caught in these areas. This assumption may over- or under-estimate catch levels for fish that have spent time in UK saltmarsh. To consider the possibility that fish caught in these areas may have used saltmarsh from other countries, we normalised the catch in each ICES area according to number of countries that share a coastal border with that area. For each species, we obtained total live weight (tonnes) for each year recorded, 2006-2016 (ICES 2018). We then calculated the average total live weight across all ten years. This average total live weight per year was then multiplied by the monetary value in GBP (£) per tonne for each species. To calculate the most recent estimation of value per tonne, we used time series data (2011-2015 MMO) from the Marine Management Organisation (MMO). Using the most recent data from 2015, we calculated the total yearly landings values (GBP) for each species landed in UK ports, which was then divided by the total amount of each species landed (tonnes), resulting in an estimate of value per tonne.

**Figure 1.**
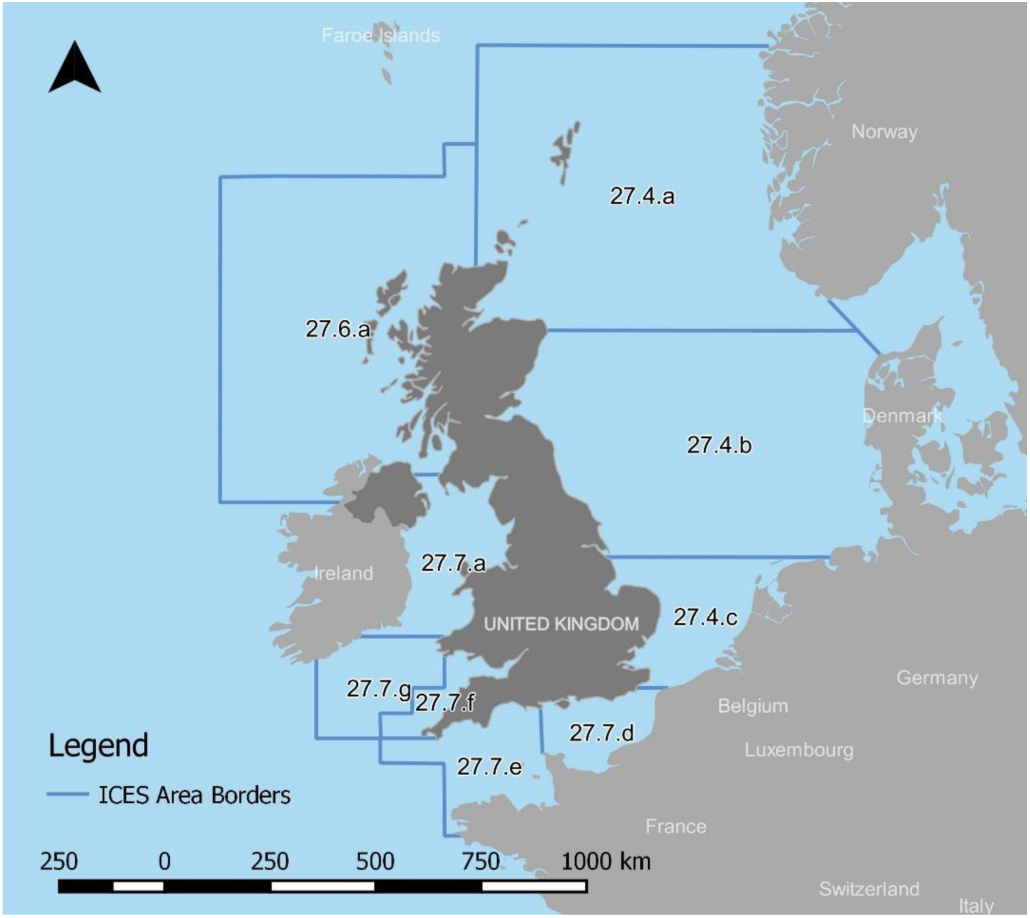
Map of the nine ICES areas that border the UK, in which fish that have spent time in UK saltmarsh are most likely to be caught. We used 2006-2016 catch data from these areas to obtain average catch per year for our target species: <u>Dicentrarchus</u> <u>labrax</u>, <u>Pleuronectes</u> <u>platessa</u>, <u>Solea</u> <u>solea</u>, <u>Chelon</u> <u>ramada</u>, and <u>Chelon</u> <u>labrosus</u>. Yearly catch in any area that borders additional countries was divided by the number of bordering countries to account for overestimation.

### Applying the Saltmarsh Residency Index

To apply the saltmarsh residency index to landings values, we multiplied the SRI value for each of our five target fish species by the commercial fisheries value for that species. We then added these together to quantify the total commercial value for saltmarsh (Jackson et al., 2015). This is shown in the following equation 4:

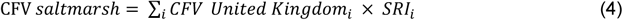

where *CFV saltmarsh* is the total commercial value of saltmarsh, *CFV United Kingdomm*_*i*_ is the commercial value for species *i*, and *SRI*_*i*_ is the saltmarsh residency index value for species *i*.

### Sensitivity Analysis

We conducted a sensitivity analysis to test the responsiveness of the final commercial value of saltmarsh to different estimates of the proportion of time adults and juveniles spend in saltmarsh. We focused on this time proportion as it is a key element of the assumption underpinning our analysis: that residency is a proxy for dependency (Scott 1999). To conduct this analysis, we calculated a species-specific sensitivity index for incremental changes in the proportion of time a species spends in saltmarsh. We used the following equation from Yeo (1991):

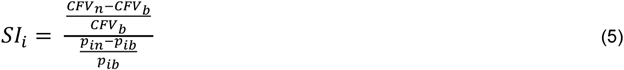

where *SI*_*i*_ is the sensitivity index for proportion of time, *P*_*ib*_ is the original total proportion of time spent in saltmarsh, *P*_*in*_ is the increased total proportion of time, *CFV*_*b*_ is the original commercial fisheries value calculated with *P*_*ib*_ and *CFV*_*n*_ is the new commercial fisheries value calculated with *P*_*in*_.

We conducted all analyses using R Version 3.5.1 (R Core Team 2018). Additional packages “plyr” (Wickham 2011),”ggplot2” (Wickham 2016) and “reshape” (Wickham 2007) were used for the economic analysis and for creating graphs.

## RESULTS

The total UK landings value for *Dicentrachus labrax, Pleuronectus platessa*, and *Solea solea* in 2015 was £19.4 million (Richardson 2017). Of this total, between £4.3 million (22.2%) and £4.8 million (24.7%) can be attributed to saltmarsh (Figure 2). The ICES catch dataset used for our analysis did not report any UK landings for either *C. ramada* or *C. labrosus*, so these species were excluded from UK totals reported above. The value of European Union commercial fleet landings for our five target species was £52.42 million in 2015 (European Commission 2018). The contribution of UK saltmarsh to this total was between £23.9 million and £25.66 million, or between 45.6% and 49% of the 2015 total. These five species are estimated to spend between 6% and 22% of their juvenile life stage in saltmarsh, where they are protected from predators and have access to safe, plentiful feeding grounds (Green et al. 2012).

**Figure 2.**
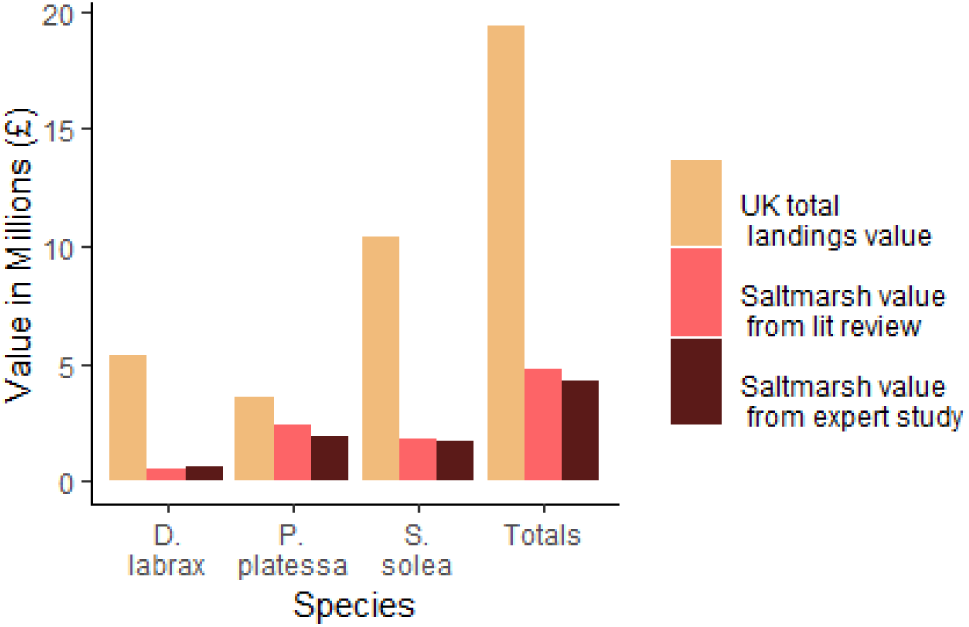
Total commercial value for each species (UK total landings value) and UK commercial fisheries value of saltmarsh as calculated with proportion of time estimates from the literature review (Saltmarsh value from lit review) and with proportion of time estimates from the expert study (Saltmarsh value from expert study) for the three focus species with recorded UK landings: <u>Dicentrarchus</u> <u>labrax</u>, <u>Pleuronectes</u> <u>platessa</u>, and <u>Solea</u> <u>solea</u>. Totalled estimates across all 3 species of saltmarsh value from UK total landings, literature review, and expert study are also displayed.

The provisioning services provided by UK saltmarsh to UK commercial fisheries were highest for *P. platessa*, with its estimated value attributed to saltmarsh ranging from £1.9 million to £2.4 million (26.5% to 33.5% of its total commercial landings value) (Figure 2). This large contribution can be attributed to two factors: 1) this species has the highest average SRI value of all 5 species (0.3, see Table 2, & Figure 2) and 2) an average yearly catch rate at least six times higher than any other species. The species with the lowest economic contribution was *D. labrax* (besides *C. labrosus* and *C. ramada*, which had no recorded UK landings). This is likely due to low catch levels in 2015 caused by catch limits set by the EU, which meant that its potential saltmarsh economic contribution was low (European Commission n.d.).

**Table 2.** For the five target species, estimates from the literature review for juvenile (x_sa_) and adult (y_a_) proportion of time in saltmarsh used to calculate (SRI_a_) (equation 1). Estimates from the expert study for juvenile (x_b_) and adult (y_b_) proportion of time in saltmarsh used to calculate (SRI_b_) (equation 1). SRI average is the mean of (SRI_a_) and (SRI_b_). Sensitivity Index (SI_i_) shows the degree to which SRI changes with regard to changes in proportion of time.

| Species Name | Scientific Name | $x_a$ | $y_a$ | $SRI_a$ | $x_b$ | $y_b$ | $SRI_b$ | $SRI$ average | $SI_i$ |
| --- | --- | --- | --- | --- | --- | --- | --- | --- | --- |
| European seabass | *Dicentrarchus labrax* | 0.125 | 0.1 | 0.203 | 0.222 | 0.088 | 0.268 | 0.236 | 0.887 |
| European plaice | *Pleuronectus platessa* | 0.07 | 0 | 0.265 | 0.051 | 0.01 | 0.335 | 0.3 | 0.857 |
| Common sole | *Solea solea* | 0.06 | 0 | 0.277 | 0.054 | 0.007 | 0.295 | 0.286 | 0.837 |
| Thicklip grey mullet | *Chelon labrosus* | 0.134 | 0.116 | 0.206 | 0.183 | 0.079 | 0.21 | 0.208 | 0.998 |
| Thinlip grey mullet | *Chelon ramada* | 0.089 | 0.116 | 0.175 | 0.239 | 0.105 | 0.266 | 0.221 | 0.904 |

The calculated saltmarsh residency index (SRI) values (Table 2) represent each species’ dependency on saltmarsh. Interestingly, this dependency does not scale linearly with the estimated amount of time each species spends in saltmarsh. For example, *D. labrax* was estimated to spend the most time in saltmarsh in both the literature review and the expert elicitation process (Figure 2). However, the average SRI value for *D. labrax* (0.236) was less than that of both *S. solea* (0.286) and *P. platessa* (0.3) (Figure 2). The average SRI for *C. ramada* was lower than that of *D. labrax* at 0.221, while the average SRI for *C. labrosus* was the lowest at 0.208 (Figure 2).

The sensitivity analysis shows that a 10% increase in both proportion of time spent as a juvenile and proportion of time spent as an adult result in sensitivity index values that range from 0.837 to 0.998 across our five target species. Sensitivity index values are bounded between 0 and 1, with 0 representing not being sensitive at all and 1 meaning maximum sensitivity of commercial fisheries value to proportion of time spent in saltmarsh. A sensitivity index value of 0.88 signifies that estimates of the fisheries value of saltmarsh are highly sensitive to proportion of time estimates. This sensitivity explains the £460,000 range between the total UK commercial fisheries value (*CFV saltmarsh*) calculated using expert estimates (£4.3 million), and the *CFV saltmarsh* calculated using estimates from the literature review (£4.8 million) (Figure 2).

## DISCUSSION

In this study, we exemplify a species-specific approach to estimate the value of saltmarsh for commercial fisheries. Our approach demonstrates how demographic and economic modelling can be effectively combined to more accurately estimate saltmarsh ecosystem services value. We found that the commercial value of UK saltmarsh to the target species, European seabass (*Dicentrarchus labrax*), European plaice (*Pleuronectes platessa*), common sole (*Solea solea*), thinlip grey mullet (*Chelon ramada*), and thicklip grey mullet (*Chelon labrosus*), when landed in the UK, ranges between £4.3 million and £4.8 million per year. This implies that 22.2% and 24.7% of the total commercial UK-landings value of these species (£19.4 million) can be attributed to saltmarsh (Richardson 2017).

A key finding from our results was that the Species Residency Index (SRI), or dependency of a species on saltmarsh for habitat, did not scale linearly with the estimated amount of time each species spends in saltmarsh. For example, results showed that the SRI for *D. labrax* (0.236), found to spend the most time in saltmarsh, was less than that of both *S. solea* (0.286) and *P. platessa* (0.3) (Figure 2). This finding suggests that a linear approach to calculating the fisheries value of saltmarsh, such as that used by Turner et al. (2007) and Luisetti et al. (2011), does not accurately estimate the economic value of this provisioning service. The same can be said of using linear approaches to calculate the fisheries value of other coastal habitats. For example, Valdez et al. (2008) used linear scaling to estimate the fisheries value of mangroves. Our results suggest that methods like these may not accurately represent a species’ habitat dependency or the true economic value of a habitat to fisheries.

Saltmarsh managed realignment has taken place in over 40 locations across the UK (Maddock 2011; Colclough 2018). The Environment Agency’s estimated costs for restoring the UK’s original saltmarsh extent is £16 million per year (DEFRA & Environment Agency 2006). According to our results, saltmarsh economic contributions to UK commercial fisheries for the target species make up between 26.9%-30% of these yearly costs. Various studies have estimated the value of additional saltmarsh ecosystem services. For example, Luisetti et al. (2011) estimated that the recreational value of saltmarsh is £621 per hectare each year at the Humber estuary managed realignment site. When this estimate is extrapolated across the entire UK saltmarsh extent, saltmarsh recreational value could be as much as £28 million. Additionally, Barbier et al. (2011) estimated that the value of saltmarsh as grazing vegetation for livestock is £15.27 per hectare ever year, which when extrapolated offers an estimate of £687,150 across the UK. Our results, together with these examples, demonstrate that saltmarsh ecosystem service benefits greatly outweigh the costs of managed realignment.

The relationship between fisheries decline and saltmarsh loss has not been studied extensively. However, the availability and connectivity of coastal foraging grounds and pelagic spawning sites has been shown to contribute to the success of demersal fish populations (Martinho et al. 2007). The UK Biodiversity Action Plan estimated that 100 hectares of UK saltmarsh are lost every year (Maddock 2011). Based on our results, there are significant opportunity costs associated with continued saltmarsh decline. Similarly, there is evidence that fish populations can show strong site fidelity to coastal feeding grounds (Doyle et al. 2017). This indicates that destruction of saltmarsh could lead to habitat fragmentation and insufficient nursery and feeding grounds (Doyle et al. 2017). In this case, our results show that ecological implications could translate into economic implications. The Northern European *D. labrax* stock has experienced declining recruitment, and ICES have implemented trawling and catch size restrictions to save the stock (López et al. 2015). Maintaining existing saltmarsh areas or undertaking managed realignment to increase saltmarsh extent may help with these measures.

In this research, we provided a robust methodology to quantify species dependence on saltmarsh, and subsequently estimate the value of saltmarsh for commercial fisheries. Our methodology could also be used to estimate species dependence on other threatened marine and coastal habitats, and their economic value for commercial fisheries. For example, mangroves are an economically and ecologically important coastal fish habitat (Ronnback 1999; Valdez et al. 2008). However, different species of fish use mangroves for varying purposes and at varying life stages (Ronnback 1999; Valdez et al. 2008). Previous studies have quantified the economic fisheries value of mangroves for fish that use mangroves during their entire lifecycle (Ronnback 1999; Valdez et al. 2008). These studies do not consider the value of fish species that use mangroves only as a nursery habitat (Ronnback 1999; Valdez et al. 2008). Using residency index methods to incorporate life-stage variation in habitat use captures the value of a habitat for all species that use it during their lifecycle. Consequently, this approach provides a more complete estimate of coastal habitats’ economic fisheries value. In the absence of available landings data, a habitat residency can be calculated using demographic and habitat use data. However, to estimate the economic contribution of the habitat in question, this index value must then be applied to relevant landings values.

The methods presented in this paper provide a flexible framework to estimate the value of saltmarsh for fisheries habitat. An area for future study would be quantifying the energy transfer from one trophic level to another, which can be done through the demographic modelling approach we developed here (Yeakel et al. 2011; Xia & Yamakawa 2018), and then calculate the economic value of the biota that spend time in saltmarsh and later become food for commercial species. Quantifying the importance of saltmarsh as a habitat for recreationally-important species has also been identified as an important area for further research (Drew Associates 2004; Brown et al. 2013). This is especially relevant for recreational *D. labrax* landings, which estimates show were between 26-49% of total commercial landings in 2012 (Brown et al. 2013). Another suggestion for future fisheries valuation studies would be to explore different ways of parameterising species’ dependence on saltmarsh. A parameter that could be used is the instantaneous feeding ratio, which compares the fullness of a fish’s stomach before entering saltmarsh with the same fish’s stomach contents upon leaving saltmarsh (Laffaille et al. 1998).

Our analysis quantifies the value of saltmarsh for fisheries by accurately representing species demography and habitat use. Past studies have calculated this value using general estimates that do not consider the unique biotic makeup of UK systems. For example, Turner et al. (2007) estimated £20 million per year could be attributed to UK saltmarsh as a habitat for fish. This study was based on estimates from US saltmarsh productivity, and its figures are an order of magnitude larger than the present, species-specific estimates (Woodward & Wui 2001; Turner et al. 2007). As sea levels continue to rise (Wolters et al. 2005) and claim coastal habitats that offer valuable ecological and economic benefits to the UK (Barbier et al. 2011b), policy makers must prioritise action for conservation (Jones et al. 2011). To efficiently and accurately allocate government resources for managed realignment projects, it is essential that both the ecological benefits, as well as the socio-economic benefits of these projects are realistically and accurately estimated. Using two different approaches, we demonstrate a region-specific, species-specific method for more accurately estimating the value of saltmarsh for fisheries. Regional-specific guidance for estimating the economic benefits of coastal habitats, as presented in our analysis, can help guide policy makers to make decisions that maximise social and ecological outcomes.

