## Supporting Info for "Using a residency index to estimate the economic value of saltmarsh provisioning services for commercially important fish species"

### SUPPORTING INFORMATION

### Appendix S1: Questionnaire

Q1 Name________________________________________________________________

Q2 Gender

- Male (1)
- Female (2)
- Other (3)
- Prefer not to say (4)

Q3 Age

- 18-35 (1)
- 35-50 (2)
- 50-65 (3)
- 65+ (4)
- Prefer not to say (5)

Q4 Affiliated Organisation(s)____________________________________________________________

Q5 Years of experience working with salt marsh

- 0-5 (1)
- 5-10 (2)
- 10-20 (3)
- 20+ (4)

Q6 Please provide your Skype name. If you do not have a Skype name, please write "none" in the space.

________________________________________________________________

Please answer the following 9 questions about European sea bass (*Dicentrarchus labrax*). In the following questions, "juvenile" refers to the life stage from birth to sexual maturity; "adult" refers to the stage from sexual maturity to death.

Q7 Of the following habitats listed, please select all of those that sea bass use as a habitat at some point in their life cycle. If there are any missing from this list, please list them at the end.

Littoral rock (1)

Littoral coarse sediment (2)

Littoral sand and muddy sand (3)

Littoral mud (4)

Littoral mixed sediments (5)

Littoral biogenic reef (6)

Coastal saltmarshes (7)

Seagrass (8)

Sublittoral sediments (9)

Pelagic water column (10)

Additional habitats (11) ________________________________________________

Q8 What is the **maximum** proportion of time **juvenile** sea bass spend in salt marsh in each of the following seasons? Please enter your responses as a value between 0 and 1.

_______ Winter (1)
_______ Spring (2)
_______ Summer (3)
_______ Autumn (4)

Q9 What is the **minimum** proportion of time **juvenile** sea bass spend in salt marsh in each of the following seasons? Please enter your responses as a value between 0 and 1.

_______ Winter (1)
_______ Spring (2)
_______ Summer (3)
_______ Autumn (4)

Q10 What is your **best estimate** of the proportion of time **juvenile** sea bass spend in salt marsh in each of the following seasons? Please enter your responses as a value between 0 and 1.

_______ Winter (1)
_______ Spring (2)
_______ Summer (3)
_______ Autumn (4)

Q11 How confident are you that the interval for each season captures the truth (between 50% (as likely as not) and 100% (absolute certainty))?

_______ Winter (1)
_______ Spring (2)
_______ Summer (3)
_______ Autumn (4)

Q12 What is the **maximum** proportion of time **adult** sea bass would spend in salt marsh in each of the following seasons? Please enter your responses as a value between 0 and 1.

_______ Winter (1)
_______ Spring (2)
_______ Summer (3)
_______ Autumn (4)

Q13 What is the **minimum** proportion of time **adult** sea bass spend in salt marsh in each of the following seasons? Please enter your responses as a value between 0 and 1.
_______ Winter (1)
_______ Spring (2)
_______ Summer (3)
_______ Autumn (4)

Q14 What is your best estimate of the proportion of time **adult** sea bass spend in salt marsh in each of the following seasons? Please enter your responses as a value between 0 and 1.
_______ Winter (1)
_______ Spring (2)
_______ Summer (3)
_______ Autumn (4)

Q15 How confident are you that the interval for each season captures the truth (between 50% (as likely as not) and 100% (absolute certainty))?
_______ Winter (1)
_______ Spring (2)
_______ Summer (3)
_______ Autumn (4)

Please answer the following 9 questions about common sole (*Solea solea*). In the following questions, "juvenile" refers to the life stage from birth to sexual maturity; "adult" refers to the stage from sexual maturity to death.

Q16 Of the following habitats listed, please select all of those that sole use as a habitat at some point in their life cycle. If there are any missing from this list, please list them at the end.

Littoral rock (1)

Littoral coarse sediment (2)

Littoral sand and muddy sand (3)

Littoral mud (4)

Littoral mixed sediments (5)

Littoral biogenic reef (6)

Coastal saltmarshes (7)

Seagrass (8)

Sublittoral sediments (9)

Pelagic water column (10)

Additional habitats (11) ________________________________________________

Q17 What is the **maximum** proportion of time **juvenile** sole spend in salt marsh in each of the following seasons? Please enter your responses as a value between 0 and 1.
_______ Winter (1)
_______ Spring (2)
_______ Summer (3)
_______ Autumn (4)

Q18 What is the **minimum** proportion of time **juvenile** sole spend in salt marsh in each of the following seasons? Please enter your responses as a value between 0 and 1.
_______ Winter (1)
_______ Spring (2)
_______ Summer (3)
_______ Autumn (4)

Q19 What is your best estimate of the proportion of time **juvenile** sole spend in salt marsh in each of the following seasons? Please enter your responses as a value between 0 and 1.
_______ Winter (1)
_______ Spring (2)
_______ Summer (3)
_______ Autumn (4)

Q20 How confident are you that the interval for each season captures the truth (between 50% (as likely as not) and 100% (absolute certainty))?
_______ Winter (1)
_______ Spring (2)
_______ Summer (3)
_______ Autumn (4)

Q21 What is the **maximum** proportion of time **adult** sole would spend in salt marsh in each of the following seasons? Please enter your responses as a value between 0 and 1.
_______ Winter (1)
_______ Spring (2)
_______ Summer (3)
_______ Autumn (4)

Q22 What is the **minimum** proportion of time **adult** sole spend in salt marsh in each of the following seasons? Please enter your responses as a value between 0 and 1.
_______ Winter (1)
_______ Spring (2)
_______ Summer (3)
_______ Autumn (4)

Q23 What is your best estimate of the proportion of time **adult** sole spend in salt marsh in each of the following seasons? Please enter your responses as a value between 0 and 1.
_______ Winter (1)
_______ Spring (2)
_______ Summer (3)
_______ Autumn (4)

Q24 How confident are you that the interval for each season captures the truth (between 50% (as likely as not) and 100% (absolute certainty))?
_______ Winter (1)
_______ Spring (2)
_______ Summer (3)
_______ Autumn (4)

Please answer the following 9 questions about plaice (*Pleuronectes platessa*). In the following questions, "juvenile" refers to the life stage from birth to sexual maturity; "adult" refers to the stage from sexual maturity to death.

Q25 Of the following habitats listed, please select all of those that plaice use as a habitat at some point in their life cycle. If there are any missing from this list, please list them at the end.

Littoral rock (1)

Littoral coarse sediment (2)

Littoral sand and muddy sand (3)

Littoral mud (4)

Littoral mixed sediments (5)

Littoral biogenic reef (6)

Coastal saltmarshes (7)

Seagrass (8)

Sublittoral sediments (9)

Pelagic water column (10)

Additional habitats (11) ________________________________________________

Q26 What is the **maximum** proportion of time **juvenile** plaice spend in salt marsh in each of the following seasons? Please enter your responses as a value between 0 and 1.

_______ Winter (1)
_______ Spring (2)
_______ Summer (3)
_______ Autumn (4)

Q27 What is the **minimum** proportion of time **juvenile** plaice spend in salt marsh in each of the following seasons? Please enter your responses as a value between 0 and 1.

_______ Winter (1)
_______ Spring (2)
_______ Summer (3)
_______ Autumn (4)

Q28 What is your best estimate of the proportion of time **juvenile** plaice spend in salt marsh in each of the following seasons? Please enter your responses as a value between 0 and 1.

_______ Winter (1)
_______ Spring (2)
_______ Summer (3)
_______ Autumn (4)

Q29 How confident are you that the interval for each season captures the truth (between 50% (as likely as not) and 100% (absolute certainty))?

_______ Winter (1)
_______ Spring (2)
_______ Summer (3)
_______ Autumn (4)

Q30 What is the **maximum** proportion of time **adult** plaice would spend in salt marsh in each of the following seasons? Please enter your responses as a value between 0 and 1.
_______ Winter (1)
_______ Spring (2)
_______ Summer (3)
_______ Autumn (4)

Q31 What is the **minimum** proportion of time **adult** plaice spend in salt marsh in each of the following seasons? Please enter your responses as a value between 0 and 1.
_______ Winter (1)
_______ Spring (2)
_______ Summer (3)
_______ Autumn (4)

Q32 What is your best estimate of the proportion of time **adult** plaice spend in salt marsh in each of the following seasons? Please enter your responses as a value between 0 and 1.
_______ Winter (1)
_______ Spring (2)
_______ Summer (3)
_______ Autumn (4)

Q33 How confident are you that the interval for each season captures the truth (between 50% (as likely as not) and 100% (absolute certainty))?
_______ Winter (1)
_______ Spring (2)
_______ Summer (3)
_______ Autumn (4)

Please answer the following 9 questions about thin-lipped grey mullet (*Chelon ramada*). In the following questions, "juvenile" refers to the life stage from birth to sexual maturity; "adult" refers to the stage from sexual maturity to death.

Q34 Of the following habitat types listed, please select all of those that thin-lipped grey mullet use as a habitat at some point in their life cycle. If there are any missing from this list, please list them at the end.

Littoral rock (1)

Littoral coarse sediment (2)

Littoral sand and muddy sand (3)

Littoral mud (4)

Littoral mixed sediments (5)

Littoral biogenic reef (6)

Coastal saltmarshes (7)

Seagrass (8)

Sublittoral sediments (9)

Pelagic water column (10)

Additional habitats (11) ________________________________________________

Q35 What is the **maximum** proportion of time **juvenile** thin-lipped grey mullet spend in salt marsh in each of the following seasons? Please enter your responses as a value between 0 and 1.
_______ Winter (1)
_______ Spring (2)
_______ Summer (3)
_______ Autumn (4)

Q36 What is the **minimum** proportion of time **juvenile** thin-lipped grey mullet spend in salt marsh in each of the following seasons? Please enter your responses as a value between 0 and 1.
_______ Winter (1)
_______ Spring (2)
_______ Summer (3)
_______ Autumn (4)

Q37 What is the your best estimate of the proportion of time **juvenile** thin-lipped grey mullet spend in salt marsh in each of the following seasons? Please enter your responses as a value between 0 and 1.
_______ Winter (1)
_______ Spring (2)
_______ Summer (3)
_______ Autumn (4)

Q38 How confident are you that the interval for each season captures the truth (between 50% (as likely as not) and 100% (absolute certainty))?
_______ Winter (1)
_______ Spring (2)
_______ Summer (3)
_______ Autumn (4)

Q39 What is the **maximum** proportion of time **adult** thin-lipped grey mullet would spend in salt marsh in each of the following seasons? Please enter your responses as a value between 0 and 1.
_______ Winter (1)
_______ Spring (2)
_______ Summer (3)
_______ Autumn (4)

Q40 What is the **minimum** proportion of time **adult** thin-lipped grey mullet spend in salt marsh in each of the following seasons? Please enter your responses as a value between 0 and 1.
_______ Winter (1)
_______ Spring (2)
_______ Summer (3)
_______ Autumn (4)

Q41 What is your best estimate of the proportion of time **adult** thin-lipped grey mullet spend in salt marsh in each of the following seasons? Please enter your responses as a value between 0 and 1.
_______ Winter (1)
_______ Spring (2)
_______ Summer (3)
_______ Autumn (4)

Q42 How confident are you that the interval for each season captures the truth (between 50% (as likely as not) and 100% (absolute certainty))?
_______ Winter (1)
_______ Spring (2)
_______ Summer (3)
_______ Autumn (4)

Please answer the following 9 questions about thick-lipped grey mullet (*Chelon labrosus*). In the following questions, "juvenile" refers to the life stage from birth to sexual maturity; "adult" refers to the stage from sexual maturity to death.

Q43 Of the following habitat types listed, please select all of those that thick-lipped grey mullet use as a habitat at some point in their life cycle. If there are any missing from this list, please list them at the end.

Littoral rock (1)

Littoral coarse sediment (2)

Littoral sand and muddy sand (3)

Littoral mud (4)

Littoral mixed sediments (5)

Littoral biogenic reef (6)

Coastal saltmarshes (7)

Seagrass (8)

Sublittoral sediments (9)

Pelagic water column (10)

Additional habitats (11) ________________________________________________

Q44 What is the **maximum** proportion of time **juvenile** thick-lipped grey mullet spend in salt marsh in each of the following seasons? Please enter your responses as a value between 0 and 1.
_______ Winter (1)
_______ Spring (2)
_______ Summer (3)
_______ Autumn (4)

Q45 What is the **minimum** proportion of time **juvenile** thick-lipped grey mullet spend in salt marsh in each of the following seasons? Please enter your responses as a value between 0 and 1.
_______ Winter (1)
_______ Spring (2)
_______ Summer (3)
_______ Autumn (4)

Q46 What is your best estimate of the proportion of time **juvenile** thick-lipped grey mullet spend in salt marsh in each of the following seasons? Please enter your responses as a value between 0 and 1.
_______ Winter (1)
_______ Spring (2)
_______ Summer (3)
_______ Autumn (4)

Q47 How confident are you that the interval for each season captures the truth (between 50% (as likely as not) and 100% (absolute certainty))?
_______ Winter (1)
_______ Spring (2)
_______ Summer (3)
_______ Autumn (4)

Q48 What is the **maximum** proportion of time **adult** thick-lipped grey mullet would spend in salt marsh in each of the following seasons? Please enter your responses as a value between 0 and 1.
_______ Winter (1)
_______ Spring (2)
_______ Summer (3)
_______ Autumn (4)

Q49 What is the **minimum** proportion of time **adult** thick-lipped grey mullet spend in salt marsh in each of the following seasons? Please enter your responses as a value between 0 and 1.
_______ Winter (1)
_______ Spring (2)
_______ Summer (3)
_______ Autumn (4)

Q50 What is your best estimate of the proportion of time **adult** thick-lipped grey mullet spend in salt marsh in each of the following seasons? Please enter your responses as a value between 0 and 1.
_______ Winter (1)
_______ Spring (2)
_______ Summer (3)
_______ Autumn (4)

Q51 How confident are you that the interval for each season captures the truth (between 50% (as likely as not) and 100% (absolute certainty))?
_______ Winter (1)
_______ Spring (2)
_______ Summer (3)
_______ Autumn (4)

**Appendix S2: Literature review results**

**Table 4:** Literature review and habitat classification results for Adult European seabass (*Dicentrarchus labrax*). A 1 in a habitat column shows that the corresponding source mentioned use of that habitat.

| Source code | Source | Stage | Low energy littoral rock | Littoral coarse sediment | Littoral sand and muddy sand | Littoral mud | Littoral mixed sediments | Littoral biogenic reef | Coastal saltmarshes and saline reedbeds | Littoral sediments dominated by aquatic angiosperms | Sublittoral coarse sediment | Sublittoral sand | Sublittoral mud | Sublittoral mixed sediments | Sublittoral macrophyte-dominated sediment | Pelagic water column |
|---|---|---|---|---|---|---|---|---|---|---|---|---|---|---|---|---|
| 1 | Doyle 2017 | Adult | | | | 1 | 1 | | | | | 1 | | | | |
| 2 | Spitz 2013 | Adult | | | | | | | | | | | | | | 1 |
| 3 | Dufour 2009 | Adult | | | | | 1 | | | | | | | | | 1 |
| 4 | Laffaille 2000 | Adult | | | | | | | | | | | | | | |
| 5 | Beraud 2018 | Adult | | | | | | | | | | | | | | 1 |
| 6 | Brehmer 2013 | Adult | | | | | | | | | | | | | | 1 |
| 7 | Colclough 2005 | Adult | | | | | | | 1 | | | | | | | |
| 8 | Fonseca 2011 | Adult | | | | | | | | | | | | | | 1 |
| 9 | Green 2009 | Adult | | | | | | | | | | | | | | |
| 10 | Green 2012 | Adult | | | | | | | | | | | | | | |
| 11 | Quere 2015 | Adult | | | | | | | | | | | | | | 1 |
| 12 | Hampel 2005 | Adult | | | | | | | 1 | | | | | | | |
| 13 | Jennings 1991 | Adult | | | | | | | | | | 1 | | | | 1 |
| 14 | Jennings 1992 | Adult | | | | | | | | | | | | | | |
| 15 | Joyeux 2017 | Adult | | | | | | | 1 | | | | | | | |
| 16 | Koutsogiannopoulou 2007 | Adult | | | | | | | | | | | | | | |
| 17 | Laffaille 1998 | Adult | | | | | | | | | | | | | | |
| 18 | Lopez 2015 | Adult | | | | | 1 | | | | | | | | | 1 |
| 19 | Malavasi 2013 | Adult | | | | | | | | | | | | | | |
| 20 | Martinho 2008 | Adult | | | | | | | | | | | | | | |
| 21 | Nunn 2016 | Adult | | | | 1 | | | 1 | | | | | | | |
| 22 | Parlier 2006 | Adult | | | | | | | | | | | | | | |
| 23 | Reis-Santos 2015 | Adult | | | | | | | | | | | | | | |
| 24 | Trancart 2016 | Adult | | | | | | | | | | | | | | |
| **TOTAL** | | | **0** | **0** | **0** | **2** | **3** | **0** | **4** | **0** | **0** | **2** | **0** | **0** | **0** | **8** |

**Table 5:** Literature review and habitat classification results for Juvenile European seabass (*Dicentrarchus labrax*). A 1 in a habitat column shows that the corresponding source mentioned use of that habitat.

| Source code | Source | Stage | Low energy littoral rock | Littoral coarse sediment | Littoral sand and muddy sand | Littoral mud | Littoral mixed sediments | Littoral biogenic reef | Coastal saltmarshes and saline reedbeds | Littoral sediments dominated by aquatic angiosperms | Sublittoral coarse sediment | Sublittoral sand | Sublittoral mud | Sublittoral mixed sediments | Sublittoral macrophyte-dominated sediment | Pelagic water column |
|---|---|---|---|---|---|---|---|---|---|---|---|---|---|---|---|---|
| 1 | Doyle 2017 | Juvenile | | | | | | | | | | | | | | |
| 2 | Spitz 2013 | Juvenile | | | | | | | | | | | | | | |
| 3 | Dufour 2009 | Juvenile | | | | | 1 | | | | | | | | | |
| 4 | Laffaille 2000 | Juvenile | | | | 1 | | | 1 | | | | | | | |
| 5 | Beraud 2018 | Juvenile | | | | | | | | | | | | | | 1 |
| 6 | Brehmer 2013 | Juvenile | | | | | 1 | | | | | | | | | 1 |
| 7 | Colclough 2005 | Juvenile | | | | | | | 1 | | | | | | | |
| 8 | Fonseca 2011 | Juvenile | | | | | | | 1 | | | | | | | 1 |
| 9 | Green 2009 | Juvenile | | | | | | | 1 | | | | | | | |
| 10 | Green 2012 | Juvenile | | | | 1 | | | 1 | | | | | | | |
| 11 | Quere 2015 | Juvenile | | | | | 1 | | | | | | | | | 1 |
| 12 | Hampel 2005 | Juvenile | | | | | | | 1 | | | | | | | |
| 13 | Jennings 1991 | Juvenile | | | | | 1 | | | | | | | | | |
| 14 | Jennings 1992 | Juvenile | | | | | | | | | | | | | | 1 |
| 15 | Joyeux 2017 | Juvenile | | | | | | | 1 | | | | | | | |
| 16 | Koutsogiannopoulou 2007 | Juvenile | | | | | | | 1 | | | | | | | |
| 17 | Laffaille 1998 | Juvenile | | | | | | | 1 | | | | | | | |
| 18 | Lopez 2015 | Juvenile | | | | | 1 | | | | | | | | | 1 |
| 19 | Malavasi 2013 | Juvenile | | | | | 1 | | | | | | | | | |
| 20 | Martinho 2008 | Juvenile | | | | 1 | | | | | | | | | | |
| 21 | Nunn 2016 | Juvenile | | | | | | | 1 | | | | | | | |
| 22 | Parlier 2006 | Juvenile | | | | 1 | | | 1 | | | | | | | |
| 23 | Reis-Santos 2015 | Juvenile | | | | | | | 1 | | | | | | | |
| 24 | Trancart 2016 | Juvenile | | | | 1 | | | 1 | | | | | | | |
| **TOTAL** | | | **0** | **0** | **0** | **5** | **6** | **0** | **13** | **0** | **0** | **0** | **0** | **0** | **0** | **6** |

**Table 6:** Literature review and habitat classification results for Adult Common Sole (*Solea solea*). A 1 in a habitat column shows that the corresponding source mentioned use of that habitat.

| Source code | Source | Stage | Low energy littoral rock | Littoral coarse sediment | Littoral sand and muddy sand | Littoral mud | Littoral mixed sediments | Littoral biogenic reef | Coastal saltmarshes and saline reedbeds | Littoral sediments dominated by aquatic angiosperms | Sublittoral coarse sediment | Sublittoral sand | Sublittoral mud | Sublittoral mixed sediments | Sublittoral macrophyte-dominated sediment | Pelagic water column |
|---|---|---|---|---|---|---|---|---|---|---|---|---|---|---|---|---|
| 1 | Amara 1007 | Adult | | | | | | | | | | | | | | |
| 2 | Archambault 1018 | Adult | | | | | | | | | | | | | | 1 |
| 3 | Cabral 1000 | Adult | | | | | | | | | | | | | | |
| 4 | Couturier 1008 | Adult | | | | | | | | | | | | | | |
| 5 | Cuveliers 1010 | Adult | | | | | | | | | | | | | | |
| 6 | Darnaude 1001 | Adult | | | | | | | | | | | | | | |
| 7 | Davoodi 1007 | Adult | | | | | | | | | | | | | | 1 |
| 8 | Degré 1006 | Adult | | | | | | | | | | | | | | |
| 9 | Dierking 1011 | Adult | | | | | | | | | | | | | | |
| 10 | Durieux 1010 | Adult | | | | | | | | | | | | | | |
| 11 | Eastwood 1003 | Adult | | | | | | | | | | | | | | |
| 11 | Eastwood 1001 | Adult | | | 1 | | | | | | | | | | | |
| 13 | Emanuela 1017 | Adult | | | | | | | | | | 1 | 1 | | | |
| 14 | Engelhard 1011 | Adult | | | | | | | | | | 1 | | | | 1 |
| 15 | Grati 1013 | Adult | | | 1 | | | | | | | 1 | 1 | | | |
| 16 | Green 1009 | Adult | | | | | | | | | | | | | | |
| 17 | Guinand 1011 | Adult | | | | | | | | | | | | | | |
| 18 | Kopp 1013 | Adult | | | | | | | | | | | | | | |
| 19 | Kostecki 1011 | Adult | | | | | | | | | | | | | | |

### Table 6 continued

| Source code | Source | Stage | Low energy littoral rock | Littoral coarse sediment | Littoral sand and muddy sand | Littoral mud | Littoral mixed sediments | Littoral biogenic reef | Coastal saltmarshes and saline reedbeds | Littoral sediments dominated by aquatic angiosperms | Sublittoral coarse sediment | Sublittoral sand | Sublittoral mud | Sublittoral mixed sediments | Sublittoral macrophyte-dominated sediment | Pelagic water column |
|---|---|---|---|---|---|---|---|---|---|---|---|---|---|---|---|---|
| 10 | Lacroix 1018 | Adult | | | | | | | | | | | | | | |
| 11 | Lacroix 1013 | Adult | | | | | | | | | | | | | | |
| 11 | Le Pape 1003 | Adult | | | | | | | | | | | | | | |
| 13 | Le Pape 1004 | Adult | | | | | | | | | | | | | | |
| 14 | Le Pape 1013 | Adult | | | | | | | | | | | | | | 1 |
| 15 | Le Pape 1007 | Adult | | | | | | | | | | | | | | |
| 16 | Le Pape 1003 - Quant | Adult | | | | | | | | | | | | | | |
| 17 | Martinho 1008 | Adult | | | | | | | | | | | | | | |
| 18 | Morat 1011 | Adult | | | | | | | | | | | | | | 1 |
| 19 | Morat 1014 | Adult | | | | | | | | | | | | | | |
| 30 | Morat 1014 - The great | Adult | | | | | | | | | | 1 | | | | 1 |
| 31 | Nicolas 1007 | Adult | | | | | | | | | | | | | | |
| 31 | Post 1017 | Adult | | | | | | | | | | | | | | |
| 33 | Primo 1013 | Adult | | | | | | | | | | | | | | |
| 34 | Rochette 1010 | Adult | | | | | | | | | | | | | | |
| 35 | Rabaut 1010 | Adult | | | | | | | | | | | | | | |
| 36 | Rogers 1991 | Adult | | | | | | | | | | | | | | |
| 37 | Vinagre 1009 | Adult | | | | | | | | | | | | | | |
| 38 | Vinagre 1008 | Adult | | | | | | | | | | | | | | |
| 39 | Vinagre 1006 | Adult | | | | | | | | | | | | | | |
| **TOTAL** | | | **0** | **0** | **0** | **0** | **0** | **0** | **0** | **0** | **0** | **1** | **0** | **0** | **0** | **3** |

**Table 7:** Literature review and habitat classification results for Juvenile Common Sole (*Solea solea*). A 1 in a habitat column shows that the corresponding source mentioned use of that habitat.

| Source code | Source | Stage | Low energy littoral rock | Littoral coarse sediment | Littoral sand and muddy sand | Littoral mud | Littoral mixed sediments | Littoral biogenic reef | Coastal saltmarshes and saline reedbeds | Littoral sediments dominated by aquatic angiosperms | Sublittoral coarse sediment | Sublittoral sand | Sublittoral mud | Sublittoral mixed sediments | Sublittoral macrophyte-dominated sediment | Pelagic water column |
|---|---|---|---|---|---|---|---|---|---|---|---|---|---|---|---|---|
| 1 | Amara 1007 | Juvenile | | | 1 | 1 | | | | | | | | | | |
| 2 | Archambault 1018 | Juvenile | | | | 1 | | | | | | | | | | |
| 3 | Cabral 1000 | Juvenile | | | 1 | | | | | | | 1 | | | | |
| 4 | Couturier 1008 | Juvenile | | | 1 | 1 | | | | | | 1 | | | | |
| 5 | Cuveliers 1010 | Juvenile | | | | 1 | | | | | | | | | | 1 |
| 6 | Darnaude 1001 | Juvenile | | | 1 | | | | | | | | | | | |
| 7 | Davoodi 1007 | Juvenile | | | | 1 | | | | | | | | | | 1 |
| 8 | Degré 1006 | Juvenile | | | | 1 | | | 1 | | | | | | | |
| 9 | Dierking 1011 | Juvenile | | | | | 1 | | | | | | | | | 1 |
| 10 | Durieux 1010 | Juvenile | | | 1 | 1 | | | | | | | | | | |
| 11 | Eastwood 1003 | Juvenile | | | 1 | 1 | 1 | | | | | | | | | |
| 11 | Eastwood 1001 | Juvenile | | | 1 | | | | | | | | | | | |
| 13 | Emanuela 1017 | Juvenile | | | | | | | | | | 1 | | | | |
| 14 | Engelhard 1011 | Juvenile | | | | | | | | | | | | | | 1 |
| 15 | Grati 1013 | Juvenile | | | | 1 | | | | | | | | | | |
| 16 | Green 1009 | Juvenile | | | | | | | 1 | | | | | | | |
| 17 | Guinand 1011 | Juvenile | | | 1 | 1 | | | | | | | | | | |
| 18 | Kopp 1013 | Juvenile | | | 1 | 1 | | | | | | | | | | |
| 19 | Kostecki 1011 | Juvenile | | | | 1 | | | 1 | | | | | | | |

#### Table 7 continued

| Source code | Source | Stage | Low energy littoral rock | Littoral coarse sediment | Littoral sand and muddy sand | Littoral mud | Littoral mixed sediments | Littoral biogenic reef | Coastal saltmarshes and saline reedbeds | Littoral sediments dominated by aquatic angiosperms | Sublittoral coarse sediment | Sublittoral sand | Sublittoral mud | Sublittoral mixed sediments | Sublittoral macrophyte-dominated sediment | Pelagic water column |
|---|---|---|---|---|---|---|---|---|---|---|---|---|---|---|---|---|
| 10 | Lacroix 1018 | Juvenile | | | | 1 | | | | | | | | | | 1 |
| 11 | Lacroix 1013 | Juvenile | | | 1 | | | | | | | | | | | |
| 11 | Le Pape 1003 | Juvenile | | | 1 | | | | | | | | | | | 1 |
| 13 | Le Pape 1004 | Juvenile | | | | 1 | | | | | | | | | | |
| 14 | Le Pape 1013 | Juvenile | | | | | | | | | | | | | | 1 |
| 15 | Le Pape 1007 | Juvenile | | | 1 | 1 | | | | | | | | | | |
| 16 | Le Pape 1003 - Quant | Juvenile | | | | 1 | | | | | | | | | | |
| 17 | Martinho 1008 | Juvenile | | | | 1 | | | | | | | | | | |
| 18 | Morat 1011 | Juvenile | | | | | 1 | | | | | | | | | |
| 19 | Morat 1014 | Juvenile | | | | | 1 | | | | | | | | | 1 |
| 30 | Morat 1014 - The great | Juvenile | | | | | 1 | | | | | | | | | 1 |
| 31 | Nicolas 1007 | Juvenile | | | 1 | 1 | | | | | | | | | | |
| 31 | Post 1017 | Juvenile | | | 1 | 1 | | | | | | | | | | |
| 33 | Primo 1013 | Juvenile | | | | 1 | | | | | | | | | | 1 |
| 34 | Rochette 1010 | Juvenile | | | | 1 | | | | | | | | | | |
| 35 | Rabaut 1010 | Juvenile | | | | | | 1 | | | | | | | | |
| 36 | Rogers 1991 | Juvenile | | | 1 | | | | | | | | | | | |
| 37 | Vinagre 1009 | Juvenile | | | | 1 | | | | | | 1 | | | | |
| 38 | Vinagre 1008 | Juvenile | | | | | | | | | | 1 | | | | |
| 39 | Vinagre 1006 | Juvenile | | | | 1 | | | | | | | | | | |
| **TOTAL** | | | **0** | **0** | **6** | **11** | **3** | **1** | **0** | **0** | **0** | **2** | **0** | **0** | **0** | **6** |

**Table 8:** Literature review and habitat classification results for Adult European Plaice (*Pleuronectes platessa*). A 1 in a habitat column shows that the corresponding source mentioned use of that habitat.

| Source code | Source | stage | Low energy littoral rock | Littoral coarse sediment | Littoral sand and muddy sand | Littoral mud | Littoral mixed sediments | Littoral biogenic reef | Coastal saltmarshes and saline reedbeds | Littoral sediments dominated by aquatic angiosperms | Sublittoral coarse sediment | Sublittoral sand | Sublittoral mud | Sublittoral mixed sediments | Sublittoral macrophyte-dominated sediment | Pelagic water column |
|---|---|---|---|---|---|---|---|---|---|---|---|---|---|---|---|---|
| 1 | Burrows 1004 | Adult | | | | | | | | | | | | | | |
| 2 | Ciotti 1014 | Adult | | | | | | | | | | | | | | |
| 3 | Raedemaecker 1011 | Adult | | | | | | | | | | | | | | |
| 4 | Raedemaecker 1011 - macrobenthic | Adult | | | | | | | | | | | | | | |
| 5 | Dutz 1016 | Adult | | | 1 | 1 | | | | | | | | | | |
| 6 | Eriksson 1005 | Adult | | | | 1 | | | | | | | | | | |
| 7 | Fox 1007 | Adult | | | | | | | | | | 1 | | | | 1 |
| 8 | Freitas 1010 | Adult | | | | | | | | | | | | | | |
| 9 | Freitas 1011 | Adult | | | | | | | | | | | | | | |
| 10 | Gibson 1997 | Adult | | | | | | | | | | 1 | 1 | | | 1 |
| 11 | Gibson 1998 | Adult | | | | | | | | | | | | | | |
| 12 | Green 1009 | Adult | | | | | | | | | | | | | | |
| 13 | Haynes 1011 | Adult | | | | | | | | | | 1 | | | | 1 |
| 14 | Hufnagl 1013 | Adult | | | | | | | | | | | | | | |
| 15 | Hyder 1998 | Adult | | | | | | | | | | | | | | |
| 16 | Lauria 1011 | Adult | | | | | | | | | 1 | | | | | |
| 17 | Le Luherne 1017 | Adult | | | | | | | | | | | | | | |
| 18 | Mariani 1011 | Adult | | | | | | | | | | | | | | |
| 19 | Nash 1999 | Adult | | | | | | | | | | | | | | |
| 20 | Nash 1011 | Adult | | | | | | | | | | | | | | |
| 21 | Nunn 1016 | Adult | | | | | | | | | | | | | | |
| 22 | Pihl 1005 | Adult | | | | | | | | | | | | | | |
| 23 | Pihl 1991 | Adult | | | | | | | | | | | 1 | | | 1 |
| 24 | Rabaut 1010 | Adult | | | | | | | | | | | | | | |
| 25 | Shucksmith 1006 | Adult | | | | | | | | | | 1 | | 1 | | |
| 26 | Trimoreau 1013 | Adult | | | | | | | | | | | | | | |
| 27 | Wennhage 1007 | Adult | | | | | | | | | | | | | | |
| 28 | Wennhage 1001 | Adult | | | | | | | | | | | | | | |
| 29 | Wennhage 1007 substratum | Adult | | | | | | | | | | 1 | 1 | | | |
| **TOTAL** | | | **0** | **0** | **1** | **2** | **0** | **0** | **0** | **0** | **1** | **5** | **3** | **1** | **0** | **4** |

**Table 9:** Literature review and habitat classification results for Juvenile European Plaice (*Pleuronectes platessa*). A 1 in a habitat column shows that the corresponding source mentioned use of that habitat.

| Source code | Source | stage | Low energy littoral rock | Littoral coarse sediment | Littoral sand and muddy sand | Littoral mud | Littoral mixed sediments | Littoral biogenic reef | Coastal saltmarshes and saline reedbeds | Littoral sediments dominated by aquatic angiosperms | Sublittoral coarse sediment | Sublittoral sand | Sublittoral mud | Sublittoral mixed sediments | Sublittoral macrophyte-dominated sediment | Pelagic water column |
|---|---|---|---|---|---|---|---|---|---|---|---|---|---|---|---|---|
| 1 | Berghahn1000 | Juvenile | | | | 1 | 1 | | | | | | | | | |
| 2 | Berghahn1995 | Juvenile | | | 1 | | 1 | | | | | | | | | |
| 3 | Burrows 1004 | Juvenile | | | 1 | | | | | | | | | | | |
| 4 | Ciotti 1014 | Juvenile | | | 1 | | | | | | | | | | | |
| 5 | Raedemaecker 1011 | Juvenile | | | 1 | | | | | | | | | | | |
| 6 | Raedemaecker 1011 - macrobenthic | Juvenile | | | 1 | | | | | | | | | | | |
| 7 | Dutz 1016 | Juvenile | | | 1 | 1 | | | | | | | | | | |
| 8 | Eriksson 1005 | Juvenile | | | | 1 | | | | | | | | | | |
| 9 | Fox 1007 | Juvenile | | | | | | | | | | | | | | 1 |
| 10 | Freitas 1010 | Juvenile | | | 1 | | | | | | | | | | | |
| 11 | Freitas 1011 | Juvenile | | | | 1 | | | | | | | | | | |
| 12 | Gibson 1997 | Juvenile | | | | | | | | | | | | | | |
| 13 | Gibson 1998 | Juvenile | | | 1 | | | | | | | | | | | |
| 14 | Green 1009 | Juvenile | | | | | | | 1 | | | | | | | |
| 15 | Haynes 1011 | Juvenile | | | 1 | | | | | | | | | | | |
| 16 | Hufnagl 1013 | Juvenile | | | 1 | 1 | | | | | | | | | | |
| 17 | Hyder 1998 | Juvenile | | | | | | | | | | | | | | 1 |
| 18 | Lauria 1011 | Juvenile | | | 1 | 1 | | | | | | | | | | |
| 19 | Le Luherne 1017 | Juvenile | | | 1 | | | | | | | | | | | |
| 20 | Mariani 1011 | Juvenile | | | 1 | 1 | | | | | | | | | | 1 |
| 21 | Nash 1999 | Juvenile | | | | | | | | | | | | | | 1 |
| 22 | Nash 1011 | Juvenile | | | | 1 | | | | | | | | | | 1 |
| 23 | Nunn 1016 | Juvenile | | | | | | | 1 | | | | | | | |
| 24 | Pihl 1005 | Juvenile | | | 1 | 1 | | | | | | | | | | |
| 25 | Pihl 1991 | Juvenile | | | 1 | | | | | | | | | | | |
| 26 | Rabaut 1010 | Juvenile | | | | | | 1 | | | | | | | | |
| 27 | Shucksmith 1006 | Juvenile | | | | | | | | | | | | | | |
| 28 | Trimoreau 1013 | Juvenile | | | 1 | 1 | | | | | | | | | | |
| 29 | Wennhage 1007 | Juvenile | | | 1 | | | | | | | | | | | |
| 30 | Wennhage 1001 | Juvenile | | | 1 | | | | | | | | | | | |
| 31 | Wennhage 1007 substratum | Juvenile | | | **1** | **1** | | | | | | | | | | |
| **TOTAL** | | | **0** | **0** | **19** | **11** | **2** | **1** | **2** | **0** | **0** | **0** | **0** | **0** | **0** | **5** |

**Table 10:** Literature review and habitat classification results for Adult thicklip grey mullet (*Chelon labrosus*). A 1 in a habitat column shows that the corresponding source mentioned use of that habitat.

| Source code | Source | stage | Low energy littoral rock | Littoral coarse sediment | Littoral sand and muddy sand | Littoral mud | Littoral mixed sediments | Littoral biogenic reef | Coastal saltmarshes and saline reedbeds | Littoral sediments dominated by aquatic angiosperms | Sublittoral coarse sediment | Sublittoral sand | Sublittoral mud | Sublittoral mixed sediments | Sublittoral macrophyte-dominated sediment | Pelagic water column |
|---|---|---|---|---|---|---|---|---|---|---|---|---|---|---|---|---|
| 1 | Boglione 1006 | Adult | | | | | | | | | | | | | | |
| 1 | Gordo 1001 | Adult | | | | | | | | | | | | | | |
| 3 | Gordoa 1009 | Adult | | | | 1 | | | | | | | | | | |
| 4 | Green 1011 | Adult | | | | | | | | | | | | | | |
| 5 | Grippa 1004 | Adult | | | | | 1 | | | | | | | | | |
| 6 | Koutsogiannopoulou 1007 | Adult | | | | | | | 1 | | | | | | | 1 |
| 7 | Schaber 1011 | Adult | | | | | | | | | | | | | | |
| 8 | Trancart 1016 | Adult | | | | | | | 1 | | | | | | | |
| 9 | Uysal 1008 | Adult | | | | | 1 | | | | | | | | | |
| 10 | Vandendriessche 1007 | Adult | | | | | | | | | | | | | | |
| **TOTAL** | | | **0** | **0** | **0** | **1** | **2** | **0** | **2** | **0** | **0** | **0** | **0** | **0** | **0** | **1** |

**Table 11:** Literature review and habitat classification results for Juvenile thicklip grey mullet (*Chelon labrosus*). A 1 in a habitat column shows that the corresponding source mentioned use of that habitat.

| Source code | Source | stage | Low energy littoral rock | Littoral coarse sediment | Littoral sand and muddy sand | Littoral mud | Littoral mixed sediments | Littoral biogenic reef | Coastal saltmarshes and saline reedbeds | Littoral sediments dominated by aquatic angiosperms | Sublittoral coarse sediment | Sublittoral sand | Sublittoral mud | Sublittoral mixed sediments | Sublittoral macrophyte-dominated sediment | Pelagic water column |
|---|---|---|---|---|---|---|---|---|---|---|---|---|---|---|---|---|
| 1 | Boglione 1006 | Juvenile | | | | | 1 | | | | | | | | | |
| 2 | Gordo 1001 | Juvenile | | | | 1 | 1 | | | | | | | | | |
| 3 | Gordoa 1009 | Juvenile | | | | 1 | | | | | | | | | | |
| 4 | Green 1011 | Juvenile | | | | | | | 1 | | | | | | | |
| 5 | Grippa 1004 | Juvenile | | | | | 1 | | | | | | | | | |
| 6 | Koutsogiannopoulou 1007 | Juvenile | | | | | | | 1 | | | | | | | |
| 7 | Schaber 1011 | Juvenile | | | | | | | | | | | | | | 1 |
| 8 | Trancart 1016 | Juvenile | | | | | | | 1 | | | | | | | |
| 9 | Uysal 1008 | Juvenile | | | | | 1 | | | | | | | | | |
| 10 | Vandendriessche 1007 | Juvenile | | | | | | | | | | | | | | 1 |
| **TOTAL** | | | **0** | **0** | **0** | **2** | **1** | **0** | **3** | **0** | **0** | **0** | **0** | **0** | **0** | **2** |

**Table 12:** Literature review and habitat classification results for Adult thicklip grey mullet (*Chelon ramada*). A 1 in a habitat column shows that the corresponding source mentioned use of that habitat.

| Source code | Source | stage | Low energy littoral rock | Littoral coarse sediment | Littoral sand and muddy sand | Littoral mud | Littoral mixed sediments | Littoral biogenic reef | Coastal saltmarshes and saline reedbeds | Littoral sediments dominated by aquatic angiosperms | Sublittoral coarse sediment | Sublittoral sand | Sublittoral mud | Sublittoral mixed sediments | Sublittoral macrophyte-dominated sediment | Pelagic water column |
|---|---|---|---|---|---|---|---|---|---|---|---|---|---|---|---|---|
| 1 | Almeida 1003 | Adult | | | | | | | 1 | | | | | | | |
| 2 | Bartulović 1007 | Adult | | | | | | | | | | | | | | 1 |
| 3 | Boglione 1006 | Adult | | | | 1 | | | | | | | | | | |
| 4 | Daverat 1011 | Adult | | | | | | | | | | | | | | 1 |
| 5 | Dias 1016 | Adult | 1 | | | | | | | | | | | | | |
| 6 | França 1011 | Adult | | | | | | | 1 | | | | | | | |
| 7 | Gordo 1001 | Adult | | | | 1 | | | | | | | | | | |
| 8 | Joyeux 1017 | Adult | | | | | | | | | | | | | | |
| 9 | Laffaille 1001 | Adult | | | | | | | 1 | | | | | | | |
| 10 | Laffaille 1998 | Adult | | | | 1 | | | 1 | | | | | | | |
| 11 | Laffaille 1001 | Adult | | | | | | | 1 | | | | | | | |
| 11 | Le Pichon 1017 | Adult | | | | 1 | | | | | | | | | | |
| 13 | Lebreton 1013 | Adult | | | | | | | 1 | | | | | | | |
| 14 | Lebreton 1011 | Adult | | | | 1 | | | 1 | | | | | | | |
| 15 | Pedro 1008 | Adult | | | | 1 | | | | | | | | | | |
| 16 | Pedro 1015 | Adult | | | | 1 | | | | | | | | | | |
| 17 | Salgado 1004 | Adult | | | | 1 | | | | | | | | | | |
| 18 | Verdiell-Cubedo 1013 | Adult | | | | | | | | | | | | | | |
| **TOTAL** | | | **1** | **0** | **0** | **8** | **0** | **0** | **7** | **0** | **0** | **0** | **0** | **0** | **0** | **2** |

**Table 13:** Literature review and habitat classification results for Juvenile thinlip grey mullet (*Chelon ramada*). A 1 in a habitat column shows that the corresponding source mentioned use of that habitat.

| Source code | Source | stage | Low energy littoral rock | Littoral coarse sediment | Littoral sand and muddy sand | Littoral mud | Littoral mixed sediments | Littoral biogenic reef | Coastal saltmarshes and saline reedbeds | Littoral sediments dominated by aquatic angiosperms | Sublittoral coarse sediment | Sublittoral sand | Sublittoral mud | Sublittoral mixed sediments | Sublittoral macrophyte-dominated sediment | Pelagic water column |
|---|---|---|---|---|---|---|---|---|---|---|---|---|---|---|---|---|
| 1 | Almeida 1003 | Juvenile | | | | | | | 1 | | | | | | | |
| 1 | Bartulović 1007 | Juvenile | | | 1 | | | | | | | | | | | 1 |
| 3 | Boglione 1006 | Juvenile | | | | 1 | 1 | | | | | | | | | |
| 4 | Daverat 1011 | Juvenile | | | | | | | | | | | | | | 1 |
| 5 | Dias 1016 | Juvenile | 1 | | | | | | | | | | | | | |
| 6 | França 1011 | Juvenile | | | | | | | 1 | | | | | | | |
| 7 | Gordo 1001 | Juvenile | | | | 1 | | | | | | | | | | |
| 8 | Joyeux 1017 | Juvenile | | | | | | | 1 | | | | | | | |
| 9 | Laffaille 1001 | Juvenile | | | | | | | 1 | | | | | | | |
| 10 | Laffaille 1998 | Juvenile | | | | 1 | | | 1 | | | | | | | |
| 11 | Laffaille 1001 | Juvenile | | | | | | | 1 | | | | | | | |
| 11 | Le Pichon 1017 | Juvenile | | | | 1 | | | | | | | | | | |
| 13 | Lebreton 1013 | Juvenile | | | | | | | 1 | | | | | | | |
| 14 | Lebreton 1011 | Juvenile | | | | 1 | | | 1 | | | | | | | |
| 15 | Pedro 1008 | Juvenile | | | | 1 | | | | | | | | | | |
| 16 | Pedro 1015 | Juvenile | | | | 1 | | | | | | | | | | |
| 17 | Salgado 1004 | Juvenile | | | | 1 | | | | | | | | | | |
| 18 | Verdiell-Cubedo 1013 | Juvenile | | | | 1 | | | 1 | | | | | | | |
| **TOTAL** | | | **1** | **0** | **1** | **9** | **1** | **0** | **9** | **0** | **0** | **0** | **0** | **0** | **0** | **2** |

**Search strings:**

We conducted the literature review using the following search string for each species:

(common name OR scientific name) AND ("habitat" OR "littoral" OR "infralittoral" OR "sublittoral" OR "circalittoral" OR "rock" OR "coarse sediment" OR "sand" OR "mud" OR "mixed sediments" OR "saltmarsh" OR "salt marsh" OR "sediment" OR "deep sea" OR "pelagic")
